# Improving microbial electrosynthesis of polyhydroxybutyrate (PHB) from CO_2_ by *Rhodopseudomonas palustris* TIE-1 using an immobilized iron complex modified cathode

**DOI:** 10.1101/214577

**Authors:** Karthikeyan Rengasamy, Tahina Onina Ranaivoarisoa, Rajesh Singh, Arpita Bose

## Abstract

Microbial electrosynthesis (MES) is a promising bioelectrochemical approach to produce biochemicals. A previous study showed that *Rhodopseudomonas palustris* TIE-1 can directly use poised electrodes as electron donors for photoautotrophic growth at cathodic potentials that avoid electrolytic H_2_ production (photoelectroautotrophy). To make TIE-1 an effective biocatalyst for MES, we need to improve its electron uptake ability and growth under photoelectroautotrophic conditions. Because TIE-1 interacts with various forms of iron while using it as a source of electrons for photoautotrophy (photoferrotrophy), we tested the ability of iron-based redox mediators to enhance direct electron uptake. Our data show that soluble iron cannot act as a redox mediator for electron uptake by TIE-1 from a cathode poised at +100mV vs. Standard Hydrogen electrode. We then tested whether an immobilized iron-based redox mediator Prussian Blue (PB) can enhance electron uptake by TIE-1. Chronoamperometry indicates that cathodic current uptake by TIE-1 increased from 1.47 ± 0.04 to 5.6 ± 0.09 µA/cm^2^ (3.8 times) and the production of the bioplastic, polyhydroxybutyrate (PHB) improved from 13.5 ± 0.2 g/L to 18.8 ± 0.5 g/L (1.4 times) on electrodes coated with PB. Overall, our data show that immobilized PB can increase direct electron uptake by TIE-1 and enhances PHB production.

## 1. Introduction

Microbial electrochemical systems (MECs) use microbes to catalyze biochemical reactions at the electrode-microbe interface [1,2]. Recent research suggests that microbial electrosynthesis (MES) is an attractive approach to compensate for fossil fuel shortage and to mitigate climate change [3]. In MES, electrically driven microorganisms (e.g., cathodophilic- or metal-oxidizing microorganisms) are used as biocatalysts to convert CO_2_ to value-added chemicals, biomass or biogas using the poised cathode potential [2–7]. Under a poised potential, extracellular electron transfer between the cathode and microbes can occur in the following ways: (1) Electron transfer (ET) through H_2_, which is externally supplied from the electrolyzer, (2) ET through cathodically produced H_2_ (self-mediated), (3) Direct ET to drive microbial CO_2_ fixation [8]. Also, the production of biochemicals from CO_2_ is directly linked to the quantity of electrical energy supplied by the cathodically poised electrode [4–8].

Acetogens and methanogens are widely employed as microbial catalysts in MES for biochemical production from CO_2_ [9–11]. Under electroautotrophic conditions, both acetogens and methanogens can perform indirect extracellular electron transfer (IET) using cathodically produced H_2_ as electron mediator (self-mediator) to generate bio-chemicals. The poised cathode potentials lower than −590 mV vs. Standard Hydrogen Electrode (SHE) (or more negative potentials) can favor IET due to the production of H_2_. H_2_ even at low quantities can act as an electron mediator between the electrode and microbes [1,12,13]. Electrodes poised at low potentials of −1500 mV vs. SHE can also undergo IET due to the production of formate, which can act as an electron mediator in MES by lithoautotrophic microorganisms (e.g., *Ralstonia eutropha)* [14]. Although high levels of electron uptake can be achieved by acetogens/methanogens, the main issue associated with using them for MES is that the process requires high energy input (i.e., a more negative potential), thus increasing the cost of the biochemicals produced using this strategy [15–18].

*Rhodopseudomonas palustris* TIE-1 is an iron-oxidizing photoautotrophic microorganism that can fix CO_2_ in the presence of light by using Fe^2+^ as a source of electrons (photoferrotrophy) [19,20]. Bose *et al.* [21] demonstrated that TIE-1 can uptake electrons (~1.5 µA/cm^2^) from a solid graphite electrode under low electrical energy input (+100 mV vs. SHE). Bose *et al*. [21] showed that light enhances current uptake by TIE-1. The low energy input requirement, the use of light, the metabolic versatility and the genetic tractability of TIE-1 represent major advantages for its use in future MES applications. However, for this, we need to improve its electron uptake ability and its growth under photoelectroautotrophic conditions. Although Bose *et al.* [21] suggested that direct electron uptake is the most likely mechanism by which TIE-1 accepts electrons from an electrode poised at +100 mV vs. SHE (based on electrochemical calculations) [21], Doud *et al.* [22] suggested that ferrous iron could act as a soluble redox mediator in these experiments. Doud *et al.* [22] suggested that TIE-1 was accepting electrons via indirect electron transfer where the electrode reduced ferric iron back to ferrous iron, thus making it available to TIE-1 to be used for photoferrotrophy. The potential used by Doud *et al.* (+20 mV vs. SHE) was however different from those reported by Bose *et al.* (+100 mV vs. SHE) [21,22]. Doud *et al.* [22] also showed that increasing light input improved electron uptake by TIE-1 in an uncoupled bioelectrochemical reactor where phototrophic oxidation of Fe(II) chelated with NTA by TIE-1 produced Fe(III)-NTA. The poised electrode (+20 mV vs. SHE) reduced this back to Fe(II)-NTA [22]. This is a reaction that can occur because the Fe(III)-NTA/Fe(II)-NTA redox couple has a reduction potential of ~ +400 mV vs. SHE at circumneutral pH [23]. In contrast to studies of indirect electron uptake reported by Doud *et al.* [22], here we wanted to test the effect of addition of unchelated Fe(II) on direct electron uptake by TIE-1 from electrodes poised at +100 mV vs. SHE as reported by Bose *et al.* [21]. Our results suggest that soluble Fe(II) cannot act as a redox mediator for electron uptake by TIE-1 and is unable to enhance cathodic current uptake at +100 mV vs. SHE. In search for a redox mediator that enhances direct electron uptake by TIE-1, we decided to use an immobilized iron-based redox mediator called Prussian Blue (PB). PB is a reversible ferrous-ferric chemical complex that we electrodeposited as a film on graphite cathodes and covered with a biocompatible chitosan layer.

The use of PB for our study was motivated by previous reports where graphite cathodes modified with Fe(III) aided oxygen reduction in microbial fuel cells by acting as a redox mediator [24,25]. Also, redox mediator modified electrodes improve electron transfer in biosensors; during electrocatalysis; in charge storage devices; and for electrochromism [26]. Among these redox mediators, Prussian Blue (PB) complex {iron(III) hexacyanoferrate} is used very commonly in electrochemical biosensors [27–29]. Interestingly, an open framework structure of PB analogues allows rapid insertion and extraction of multivalent cations. PB is used as a low-cost cathode material (<$1 per Kg) in microbial batteries due to its reversible characteristic for long-term applications [28].

Here, we report that TIE-1 can accept more electrons from cathodes coated with PB, representing an inexpensive method for increasing electron uptake and the production of the bioplastic, polyhydroxybutyrate (PHB), as a product of microbial electrosynthesis. We performed electrochemical analyses to measure current uptake, electrochemical activity, and electron or charge transfer resistance across the electrode-microbe (TIE-1) interface of the unmodified and modified electrodes during photoelectroautotrophic growth. The results show that extracellular electron uptake of TIE-1 increased up to 3.8 times in the presence of the immobilized ET redox mediator, Prussian Blue.

## 2. Experimental

### 2.1. Inoculum and Bioelectrochemical cell (BEC) setup

Electron uptake (EU) experiments with TIE-1 were carried out in a seal-type single chamber electrochemical cell (C001 Seal Electrolytic cell, Xi’an Yima Opto-electrical Technology Com., Ltd, China). 10 mL of cells pre-grown in Freshwater (FW) [30] medium containing H_2_ as an electron donor and 22 mM sodium bicarbonate were inoculated in 70 mL of FW medium to achieve a final OD_660_ of ~0.01. This was followed by gas exchange for 20 mins with N_2_/CO_2_ (80%:20%), and the final headspace pressure was set as 7 psi. All photoelectroautotrophic growth experiments were replicated (n=3) at 26 °C under continuous infrared light (illumination) unless noted otherwise (Fig. S1). We performed two sets of experiments: 1) Those with the addition of soluble Fe(II) using unmodified graphite electrodes, and 2) Those using the electrode modified with PB.

### 2.1. Bioelectrochemical experiments with dissolved Fe(II)

Electron uptake of TIE-1 was performed by the addition of dissolved Fe(II) with poised electrodes in the seal type electrochemical cell. Here, **s**pectroscopically pure graphite rods (GR, 5.149 cm^2^, SPI supplies, USA) served as the working electrode, Pt foil as the counter electrode and Ag/AgCl as the reference electrode. All potential values are reported with respect to the Standard Hydrogen Electrode potential (SHE) unless specified otherwise. All electrochemical experiments were carried out using the Gamry electrochemical workstation (Gamry Multichannel potentiostat, USA). To investigate the dissolved Fe(II) during the EU experiment, about 5-6 mM of FeCl_2_ was added to FW medium in the presence or absence of TIE-1. EU by TIE-1 was measured in terms of current by chronoamperometry (CA) method at a poised potential of +100 mV vs. SHE for 152 h. Cyclic voltammetry (CV) characteristics of initial (0 h) and final (152 h) FW medium was analyzed to understand the effect of Fe(II) addition. Further, a colorimetric Ferrozine based assay was used to determine Fe(II) oxidation in the bioreactor as reported previously [20]. Finally, both the electrode surface and the spent salt medium containing planktonic cells was analyzed by JEOL JSM-7001 LVF field emission scanning electron microscopy (FE-SEM). In which, a piece (5 mm) of the graphite cathode or the spent medium from the bioreactors was fixed in 2% glutaraldehyde in 100 mM sodium cacodylate buffer for 5 h. Fixed graphite cathodes were gently rinsed with 100 mM cacodylate buffer followed by dehydration washing with a series of ethanol for 10 mins (30, 50, 70 and 100%). Finally, the dehydrated microbial cathode samples were sputter coated with a thin gold layer to perform SEM imaging and Electron Dispersive Spectroscopy (EDS).

### 2.2. Electrochemical modification of graphite cathodes

Graphite rods (GR, 5.149 cm^2^) were used as substrate electrodes for Prussian blue (PB, Fe_4_[Fe(CN)_6_]_3_ • *x*H_2_O) electrodeposition in a three electrode configured electrochemical cell as described above. Electrochemical deposition was performed using a bath containing 10 mM K_3_[Fe(CN)_6_], 10 mM FeCl_3_.6H_2_O and 10 mM HCl (Fig. S1). Electrodeposition of PB was carried out at a constant potential of −300 mV for 180 seconds using the Gamry electrochemical workstation. The modified graphite electrodes were cycled in 0.1 M KCl >30 times to get a stable reversible redox couple. Further, the PB-graphite electrodes were dip-coated with 0.5% chitosan solution and dried under N_2_ gas. Prior to use, the PB- chitosan (PB/Chit) coated graphite electrodes were immersed in deionized water for 4 h and gently rinsed to remove soluble ions on the electrode surface. In order to compare the effect of the PB modification on electron uptake by TIE-1, the working electrode was configured as an unmodified graphite rod (GR), a graphite rod coated with 0.5% chitosan (GR/Chit), and a graphite rod modified with PB and 0.5% chitosan (GR/PB/Chit). PB modified electrodes with no chitosan were not tested because of the possible detachment of the PB film in the bioreactor. Surface analysis of as-deposited PB complex was confirmed with SEM, X-ray photoelectron spectroscopy (Physical Electronics^®^ 5000 VersaProbe II Scanning ESCA (XPS) Microprobe), and the thickness of PB layer was measured with a profilometer (KLA - Tencor Alpha - Step D - 100 Profilometer).

### 2.3. Bioelectrochemical experiment with modified electrodes

To measure the current response, Chronoamperometry (CA) analysis was conducted for 130 h at the constant applied potential of +100 mV. Cyclic voltammetry (CV) and differential pulse voltammetry (DPV) of TIE-1 on different graphite electrodes was performed using a potential scan from −100 mV to +900 mV. Electrochemical impedance spectroscopy (EIS) was performed at +100 mV in the frequency range of 1 MHz to 10 mHz with a perturbation voltage of 10 mV. Potentiodynamic polarization (tafel) of electrodes with biofilms were performed from −250 mV (cathodic reduction) to + 250 mV (anodic oxidation) at 0.5 mV/s. Field Emission-SEM was used to characterize the microbial attachment on graphite cathodes, and the sample preparation for imaging was as described above.

### 2.4. PHB analysis

10 mL of TIE-1 grown under photoelectroautotrophy condition using unmodified (GR) and PB complex modified electrode (GR/PB/Chit) were pelleted at 5000 rpm and stored at - 80˚C until PHB extraction and analysis were performed. 1 mL of ultrapure LC-MS grade water and 600 µL of methanol (HPLC grade) was added to arrest metabolic activity. Samples were dried for 6 hours under vacuum in a Savant SC210A Speedvac concentrator (Thermo Fisher Scientific). 425 µL of methanol and 500 µL of HPLC grade chloroform, 75 µL of 95-98% sulfuric acid were added to the dried sample. Samples were digested for 1 hour in a water bath at 95°C. Digested samples were cooled on ice rapidly. 0.5 mL of LC-MS grade water was then added. Samples were vortexed and centrifuged at 5000 X *g* for 10 min. The organic phases were transferred into glass vials and dried for 30 min under speed vacuum. Finally, the dried samples were re-suspended in 500 µL of 50% acetonitrile + 50% water. Samples were filtered to remove any cell debris and run on Agilent Technologies 6420 Triple Quad LC/MS as follows: stationary phase was a Hypercarb column, particle 5µm, 100×2.1mm (Thermo Scientific, USA); phase A was water, 0.1% formic acid; phase B was Acetonitrile, 1% formic acid; injection volume was 5 µL; flow rate was 500 µL/min; column temperature was 15°C; gas temperature was 300°C. PHB were detected as crotonic acid with an m/z=87 [31].

## 3. Results and discussion

### 3.1. Electrotrophic characteristics of TIE-1 in response to addition of soluble Fe(II)

To understand the effect of soluble Fe(II) on electron uptake from poised electrodes (solid electron donor) by TIE-1, an unmodified graphite cathode was poised at +100 mV vs. SHE in presence of 5-6 mM soluble FeCl_2_ under both abiotic (No cells) and biotic conditions (TIE-1 cells) for 152 h. On addition of Fe(II), the cathodic current changed to anodic current (Fig. 1a) suggesting that Fe(II) is donating electrons to the cathode; in effect getting oxidized to Fe(III). The peak oxidative current was noted for the abiotic (FeCl_2_ only) and biotic systems (FeCl_2_ + TIE-1) as 21.18 ± 2.2 µA/cm^2^ (total current, 3.498 ± 0.28 mA h) and 17.61 ± 1.3 µA/cm^2^ (2.594 ± 0.11 mA h), respectively (Fig. 1a,b).

**Fig. 1.**
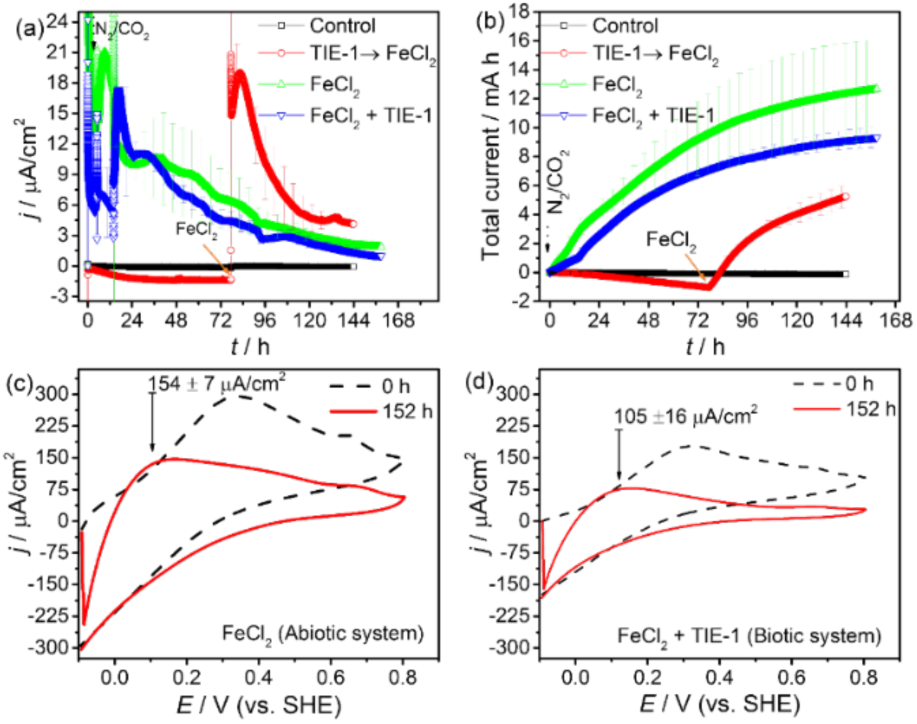
Effect of FeCl_2_ or Fe(II) dissolved freshwater (FW) medium on Electron Uptake (EU) using unmodified graphite cathodes. Chronoamperometry (a) and the total current capacity (b) of sterile (control) and TIE-1 (biotic) on an unmodified graphite electrode at a poised potential of +100mV vs. SHE for 152 h under N_2_/CO_2_. Standard deviation of replicated data (n=3) is included. Cyclic voltammetry (5 mV/s) characteristics of added Fe(II) in the abiotic (c) and biotic (d) system at the end of EU experiment.

This effect was observed from the cyclic voltammetry of both the abiotic and biotic systems (Fig. 1c,d) at the end chronoamperometry. In the abiotic system, the change in oxidation current was 154 ± 7 µA/cm^2^ compared to the biotic system (105 ± 16 µA/cm^2^) at the interval of 0 - 152 h. Overall, the presence of TIE-1 cells lowers the observed anodic current. This confirms that the electrode mediates Fe(II) oxidation. This effect is perhaps due to the continued ability of TIE-1 cells to directly uptake electrons from the poised cathode, thus competing for the electrode surface. Chronoamperometry on biotic graphite electrodes with no added Fe(II) confirms that TIE-1 accepts electrons from unmodified electrodes as reported previously (-1.39 ± 0.02 µA/cm^2^; Fig. 1a) [21]. SEM – EDS on both the abiotic and biotic reactor electrode surface and the spent medium showed the presence of iron oxides similar to Ferrihydrite (Fig. 2, Fig. 3, Fig. 4, Fig. S2). In the biotic system, the competition between Fe(II) and TIE-1 for the electrode surface was corroborated by SEM imaging of BECs, where TIE-1 cells were exposed to both a poised cathode and Fe(II) (Fig. 2a-b). SEM images show that TIE-1 cells attach to areas devoid of iron oxides (Fig. 2b-b’ and Fig 3a-b). In a parallel experiment, we grew TIE-1 cells in poised reactors for 77 h before Fe(II) addition. SEM images of these electrodes show that cells already attached to the graphite electrodes get coated with iron oxides post Fe(II) addition (Fig. 2a,a’). Overall these data suggest that Fe(II) gets oxidized by an electrode poised at +100 mV vs. SHE. These data also indicate that TIE-1 and Fe(II) compete for the electrode surface for access to electrons.

**Fig. 2.**
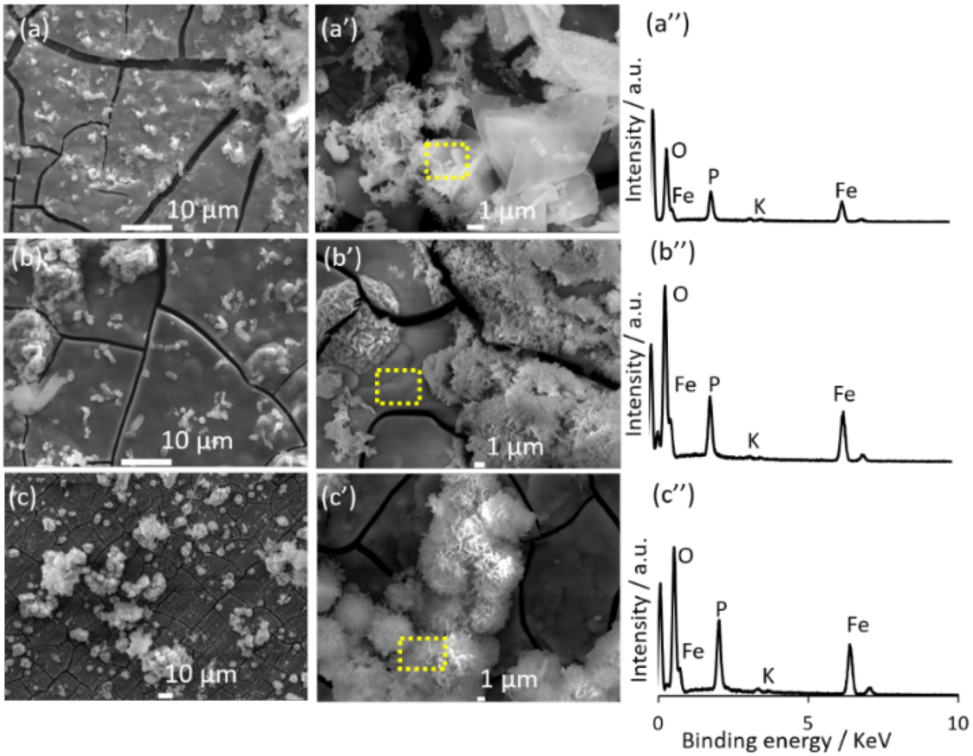
SEM images of graphite cathode at the end of the EU experiment with dissolved Fe(II) in FW medium. (a, a’) Biotic system (FeCl_2_ →TIE-1), (b, b’) biotic system (FeCl_2_ + TIE-1), and (c,c’) abiotic system (FeCl_2_). EDS (Electron Dispersive Spectroscopy) of square portion was inserted to their respective SEM images.

**Fig. 3.**
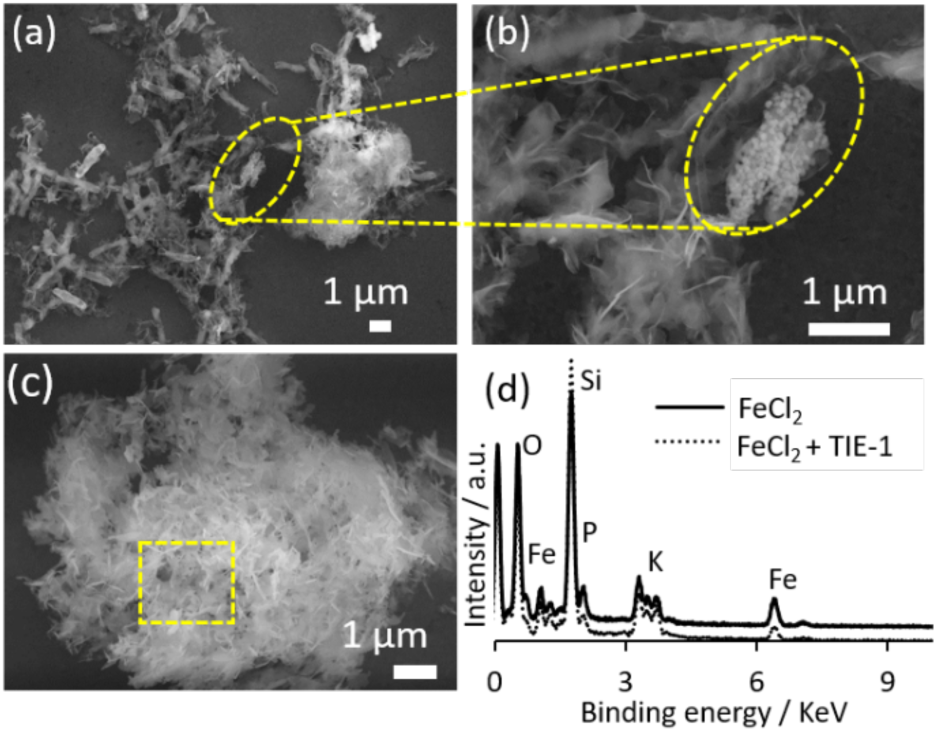
(a,b) SEM images of spent medium containing planktonic cells coated with amorphous Ferrihydrite in the biotic system (FeCl_2_ + TIE-1) and (c) abiotic system (FeCl_2_). (d) EDS (Electron Dispersive Spectroscopy) of circled portion was inserted.

**Fig. 4.**
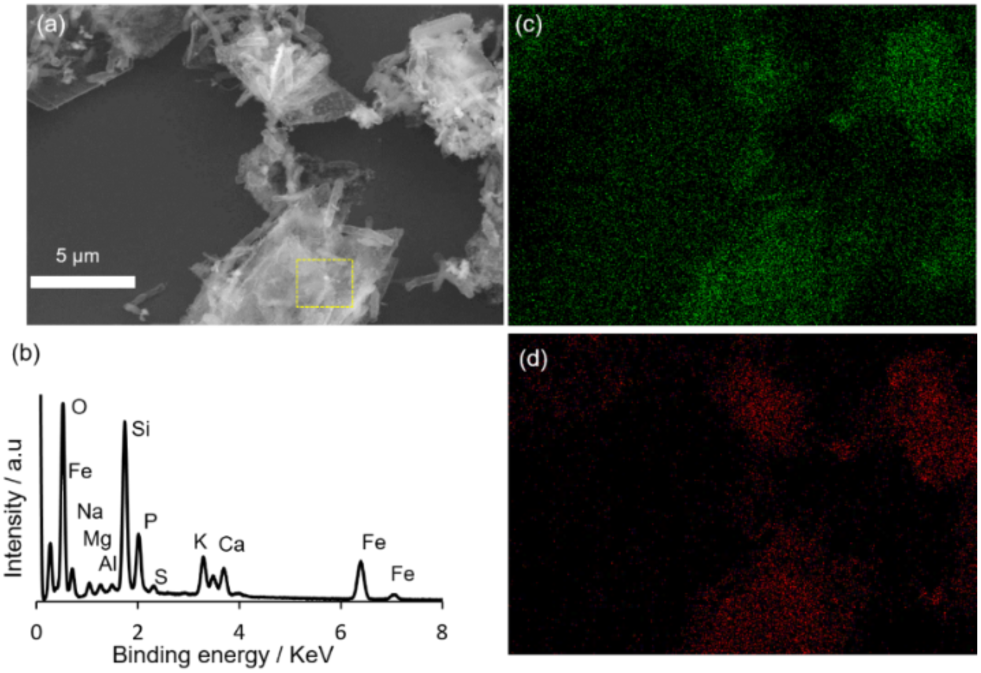
SEM image of spent medium containing planktonic cells with sheet like Ferrihydrite formation in dissolved Fe(II) reactor (FeCl_2_→TIE-1, biotic system) (a), EDS spectrum corresponds to the yellow square area (b), Elemental map of Oxygen (c), and Iron (d).

Ferrozine assays on abiotic and biotic reactors show that Fe(II) gets oxidized to Fe(III) in both cases (Table S1). In biotic reactors, 36% of the added Fe(II) is oxidized while in the abiotic reactors 19% of the Fe(II) to Fe(III). The higher Fe(II) oxidation in the biotic reactor is due to the concurrent effects of photoferrotrophy and abiotic Fe(II) oxidation by the electrodes. It’s notable that complete Fe(II) oxidation is not observed in the biotic reactors even after 152 h of incubation suggesting that TIE-1 is using both the electrodes and Fe(II) for electrons. These data clearly show that added Fe(II) cannot serve as a redox mediator to enhance cathodic electron uptake by TIE-1. In fact, Fe(II) competes with TIE-1 for the electrode surface as a source of electrons.

### 3.2. Characterization of Prussian Blue complex on graphite in abiotic systems

Cyclic voltammetry was used to characterize the electrochemical activity of the PB modified graphite cathode. Fig. 5a shows the scan rate dependent cyclic voltammetry behavior of PB with the typical characteristics of their redox peak pairs and agrees with reported results [32]. A redox peak center located at 0.42 V is due to the electrochemical transformation of PB to Prussian White (PW), while a redox peak at 1.07 V corresponds to the transformation of PG (Prussian Green) to PB. The related electrochemical reaction occurs due to electron transfer between the Fe^2+^ and Fe^3+^ complex as shown below (eqn. 1, 2) [32,33].

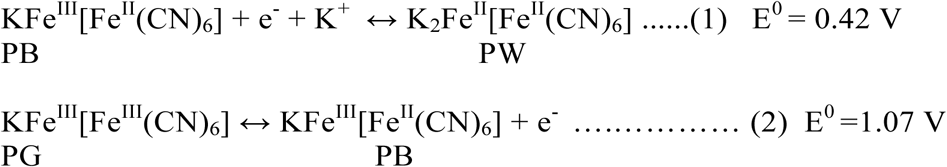

**Fig. 5.**
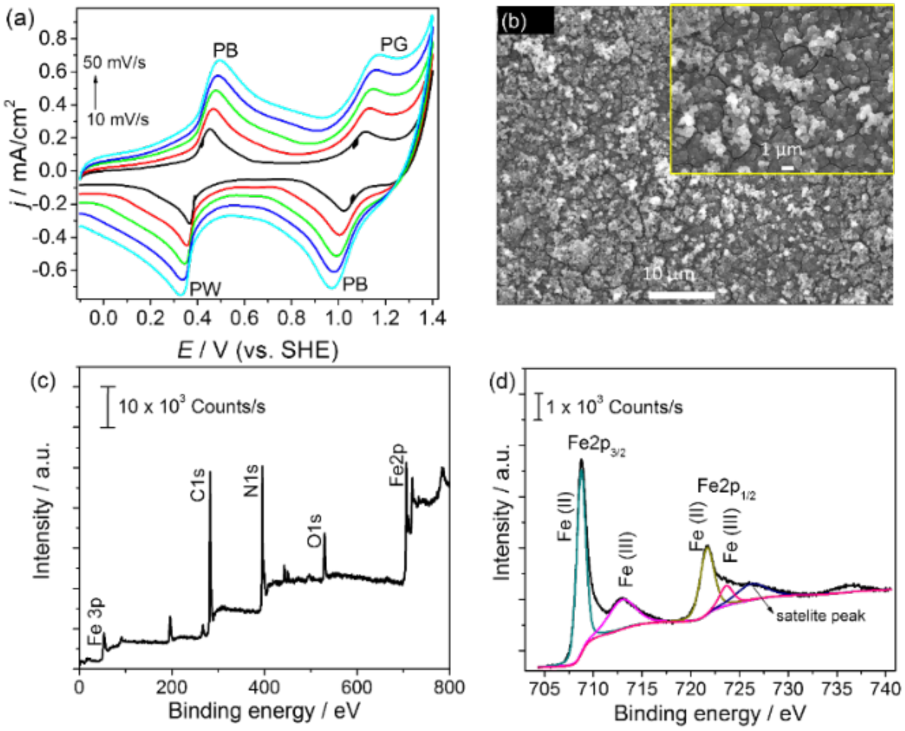
(a) Cyclic voltammetry of redox complex (PB) deposited graphite electrode in 0.1 M KCl at different scan rate; PB - Prussian blue, PW - Prussian white, PG - Prussian green, (b) SEM image of PB on graphite (insert: lower scale image), (c) X-ray photoelectron spectroscopy (XPS) of PB complex, and (d) Fe 2p XPS of PB complex.

The anodic and cathodic peak current ratio (*I*_*c*_/*I*_*a*_) of each of the redox peak potentials centered at 0.42 V and 1.07 V were 1.15 ± 0.02 and 1.01 ± 0.01, respectively. A value close to 1 indicates that the PB modified electrodes demonstrate an electrochemical redox reaction that is reversible at the graphite electrode surface [33]. Also, the peak potential differences (*∆E*) of each redox pair was in the range of 87 mV to 142 mV, which is in agreement with previous studies [34]. The plot (data not shown) of peak current (anodic and cathodic) linearly increased with the square root of the scan rate (0.997 correlation coefficient). This indicates that the electrochemical process is controlled by diffusion. The reversibility of PB (Fe^3+^ polynuclear complex) and PW (Fe^2+^ polynuclear complex) is an important characteristic for using these chemicals as redox mediators for TIE-1. It should be noted that the reduced form of PW (Fe^2+^ complex) will occur at the potential ≤0.4 vs. SHE, which could act as an electron donor for TIE-1 and accelerate extracellular electron transfer. The electrochemical deposition strategy of Prussian blue (PB) complex is well characterized in biosensor applications [35–37]. In addition, here we characterized the deposited PB on graphite using surface analytical techniques such as Scanning Electron Microscopy (SEM) with Electron Dispersive Spectroscopy (EDS), X-Ray Photon Spectroscopy (XPS), and thickness measurements using SEM and a profilometer.

### 3.3. Surface analysis of the PB complex

The electrochemically deposited PB complex was further studied using surface analytical techniques such as SEM and XPS (Fig. 5b and Fig. 5c,d). The structure of the PB matrix and elemental composition could potentially influence biofilm formation and microbial electroactivity. Fig. 5b shows the SEM of PB deposited on graphite. We saw that PB was deposited as nanoparticles with a size of 70-130 nm and formed a layer (thickness of 670- 729 nm). These PB nanoparticles can maximize the contact of microbes with the graphite surface. Further, to confirm the elemental composition of the electrodeposited PB, XPS analysis was performed. Fig. 5c shows the full range XPS spectrum of the PB complex, which consists of main peaks such as N 1s, C 1s and Fe 2p that can be clearly seen.

Also, deconvoluted XPS spectra for Fe 2p (Fig. 5d) indicate the oxidation states of Fe in the PB complex. We observe that Fe 2p is composed of two groups of peaks namely, Fe 2p_3/2_ (at a lower binding energy) and Fe 2p_1/2_ (at higher binding energy). The peaks at 708.8 eV (Fe2p_3/2_) and 721.7 eV (Fe2p_1/2_) can be correlated to the presence of Fe(II). The peaks at 713.1 eV (Fe2p_3/2_) and 723.7 eV (Fe2p_1/2_) can be assigned to Fe(III). Based on the results obtained from XPS, the electrodeposited complex can be reasonably assigned as insoluble PB complex with a formula of PB as Fe_4_^III^[Fe^II^(CN)_6_]_3_ [38–40].

### 3.4. Cathodic current uptake by TIE-1 with PB modified electrodes

The redox reversibility of the PB complex modified electrode was confirmed with CV analysis prior to use in bioelectrochemical studies (Fig. 6a). After inoculating TIE-1 in the bioreactor, the cathodic current was measured with unmodified graphite (GR-TIE-1), graphite with chitosan (GR/Chit-TIE-1) and graphite with PB/chitosan (GR-PB/Chit-TIE-1) electrode (Fig. 6). In all cases, the “no cell” control reactor did not show any significant current uptake over the operation period. The TIE-1 inoculated systems showed their ability of cathodic current uptake within 24 h, with a steep increase in cathodic current from 5 h - 48 h in all biocathodes. The maximum cathodic current (*I*_*max*_) uptake by TIE-1 was 5.6 ± 0.09 µA/cm^2^ (GR/PB/Chit-TIE-1) > 1.61 ± 0.15 µA/cm^2^ (GR/Chit-TIE-1) > 1.47 ± 0.04 µA/cm^2^ (GR-TIE-1). This indicates that the chitosan-modification alone only slightly improved current consumption compared with unmodified graphite. However, the PB modified electrode significantly enhanced the electron uptake by TIE-1 (up to 3.8 times). This effect was comparable with the cathodic electron uptake by *E. coli* using cathodes modified with cytocompatible electron mediators composed of redox polymers (7.8 µA/cm^2^) [41]. The total quantity of current consumption (Fig. 6d-e) was assessed as −1.74 ± 0.03 mA h (for GR/PB/Chit-TIE-1), which is ~3.2 times higher than the unmodified (-0.53 ± 0.01 mA h) and the chitosan modified graphite cathode (-0.61 ± 0.05 mA h). Also, the number of electrons supplied to the system was derived from the quantity of current transferred to the bioreactors (1 Faraday charge per mole electrons). It was calculated as 65.08 µmole e^−^ (GR/PB/Chit-TIE-1) > 22.90 µmole e^−^ (GR/Chit-TIE-1) > 19.89 µmole e^−^ (GR-TIE-1). The chronoamperometry data suggests that the current consumption level (high or low) can influence the biomass production rate [42]. Thus, the moles of electrons supplied to the MES reactor and moles of electrons recovered by cell biomass can be compared. By applying the molecular formula of cell biomass as CH_2.08_O_0.53_N_0.24_, 4.3 mole e^−^ are required for 1 mole of cell biomass [42]. The moles of biomass were derived as 15.12 µmol for (GR/PB/Chit-TIE-1), 5.33 µmol for (GR/Chit-TIE-1), and 4.63 µmol for (GR- TIE-1). The observed planktonic OD_660_ supports this trend; 0.023 (GR/PB/Chit-TIE-1), 0.014 (GR/Chit-TIE-1) and 0.014 (GR-TIE-1). This effect was also in agreement with the increased production of PHB from GR/PB/Chit-TIE-1 (18.8 ±0.5 g/L) and GR/Chit-TIE-1 or GR-TIE-1(13.5±0.2 g/L).

**Fig. 6.**
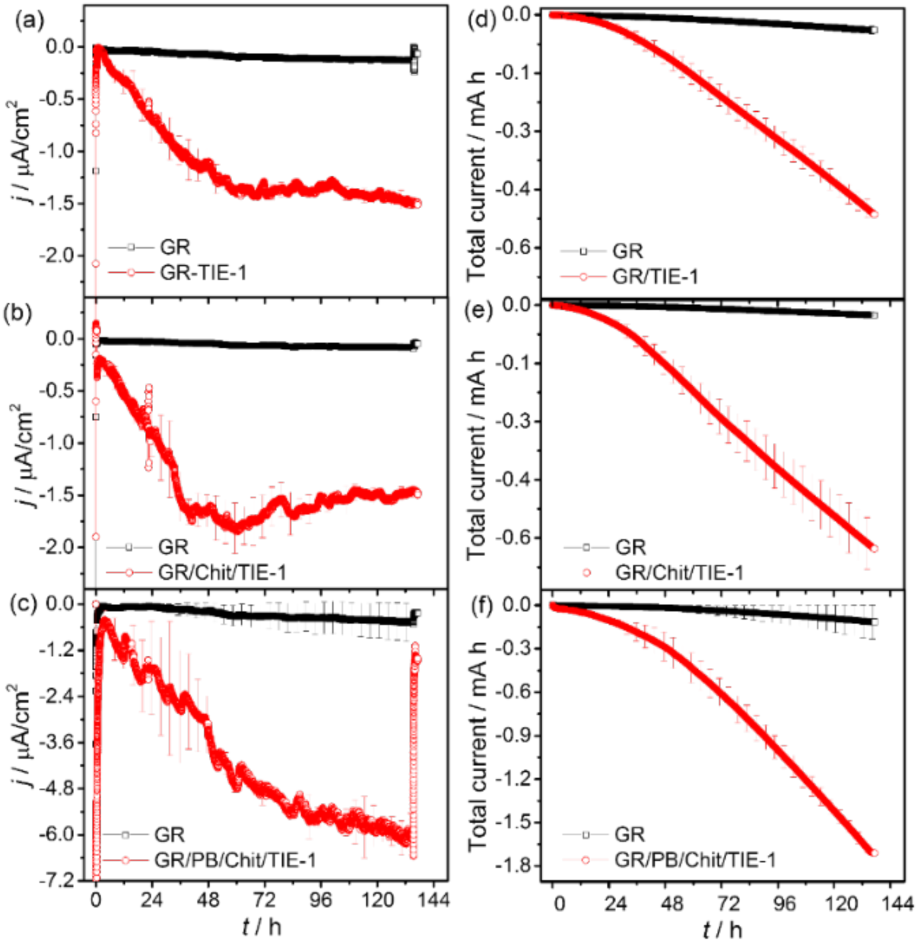
Chronoamperometry of sterile (control) and TIE-1 on a graphite electrode at poised potential of +100mV vs. SHE for 130 h under N_2_/CO_2_. Standard deviation of replicated data (n=3) were included for Current density vs. Time (a,b,c) and Total current vs. Time (d,e,f).

### 3.5. Voltammetric characteristics of the biocathodes

CV and DPV were performed to characterize the bioelectrochemical redox activity of TIE-1. Fig. 7a-c shows the CV of modified and unmodified biocathodes compared with sterile cathodes at a scan rate of 5 mV/s. The midpoint redox potentials of GR-TIE-1, GR/Chit-TIE-1 or GR/PB/Chit-TIE-1 (Fig. 7a-c) are 0.190 V, and 0.290 V, which is closely related to the redox peak potential reported previously for TIE-1 [21]. Interestingly, the PB complex modified biocathode (Fig. 7c) retains the two midpoint redox potentials of TIE-1 at 0.190 V and 0.290 V. The improved redox current was observed at 0.290 V due to the reversibility of PB (Fig. 7c). Although CV is an essential characterization technique to detect redox reactions that occur at the electrode surface, it has a low detection limit [43–45]. Pulse voltammetry techniques have frequently been used as complementary methods to CV. For pulse voltammetry techniques, the charging current can be lowered, and this lends higher sensitivity to our ability to measure Faradaic current at the redox signal [46,47]. Differential peak current (*∆I*) at the redox signal was derived from the differential pulse voltammogram (Fig. 7d-f) to measure the biofilm’s electroactivity. In all biocathodes, DPV consistently exhibits redox signals (*Ep*) with the redox potential of 0.190 V and 0.290 V as seen in the CV results. Also, the redox signal (*Ep*) at 0.290 V shows the peak differential current (*∆I*) of 88.4 µA/cm^2^ for the GR/Chit-TIE-1 cathode. This is 5.9 times higher than the unmodified biocathode (11.4 µA/cm^2^, GR-TIE-1), and is 7.6 times higher than chitosan modified biocathode (14.8 µA/cm^2^, GR/Chit-TIE-1).

**Fig. 7.**
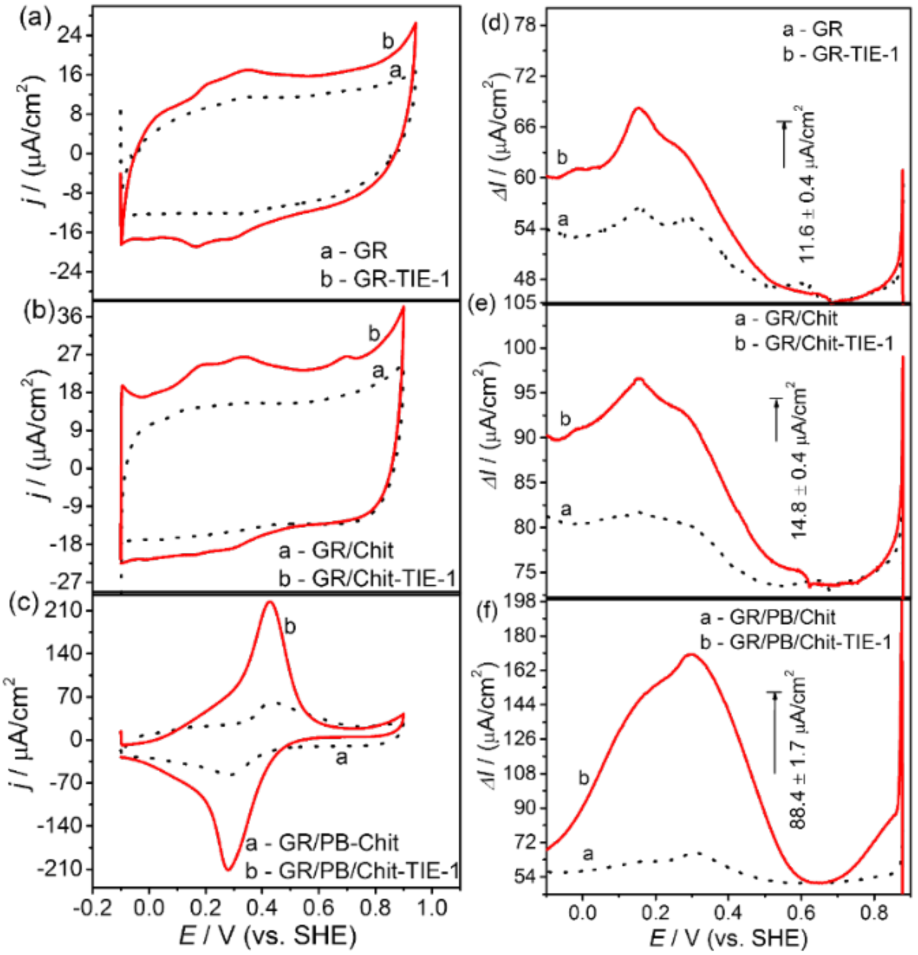
Representative cyclic voltammetry (a, b, c) of control (sterile) graphite cathodes and TIE-1 were recorded in FW medium at a scan rate of 5 mV/s under N_2_/CO_2_; Differential pulse voltammetry (Potential vs. Differential current, *∆I*) of TIE-1 on different graphite cathodes (d, e, f).

Further, the DPV results support that the PB complex modified cathode acts as an immobilized electron transfer mediator for the redox reaction at the redox peak potential of 0.290 V. The redox peak current is directly proportional to the concentration of electrochemically active molecules at the surface of the cathode. The surface covered electroactive sites in the biocathodes were calculated from the CV results by integrating charge under either the anodic or cathodic peaks. The surface coverage of electroactive moieties per unit area of the biocathode was 2.360 × 10^−10^ mol/cm^2^ for GR/PB/Chit-TIE-1, 3.0624 × 10^−11^ mol/cm^2^ for GR/Chit-TIE-1 and 1.7923 × 10 ^−11^ mol/cm^2^ for GR-TIE-1 respectively [48,49]. Based on the surface coverage value, the PB complex modified biocathode promotes the electroactivity of the biofilm by one order of magnitude compared to the unmodified and chitosan modified biocathodes per unit area. It should be further noted that chitosan has positively charged terminal groups, which may help enhance the surface functionality by attracting bacteria, and providing a microenvironment for biological reactions at the biocathode [46,50,51]. The biocathodes were scanned from 0.1 V to 0.6 V at different sweep rates from 1 to 5 mV/s in a cell-free medium solution (Fig. 8a-c). We observed that the mid-point potential of all biocathodes with TIE-1 retained their midpoint redox potentials as seen in the previous CVs of the bioreactors. A slight redox potential shift (15-20 mV) was observable perhaps due to the addition of fresh medium. The biocathode peak currents (anodic or cathodic) increased linearly with an increase in sweep rate (Fig. 8d). The linear correlation (R^2^= 0.999) of peak currents with the sweep rate indicates that the biocathode used a surface or diffusion controlled bioelectrochemical reaction [49,52]. This might be due to the effect of the electron transfer mediator at the bio-interface. Further, the supernatant of all reactors (spent medium) were analyzed for any dissolved mediator (e.g., PB complex) using voltammetry techniques such as CV (Fig. 8e) and DPV (Fig. 8f) with glassy carbon as working electrode. The results reveal that no obvious redox peaks were found in the potential region of 0.2 to 0.3 V, unlike the PB complex modified bio-cathode. This confirms that the PB complex modified biocathode does not shed PB during the experiments and that PB is surface confined when covered by a chitosan layer.

**Fig. 8.**
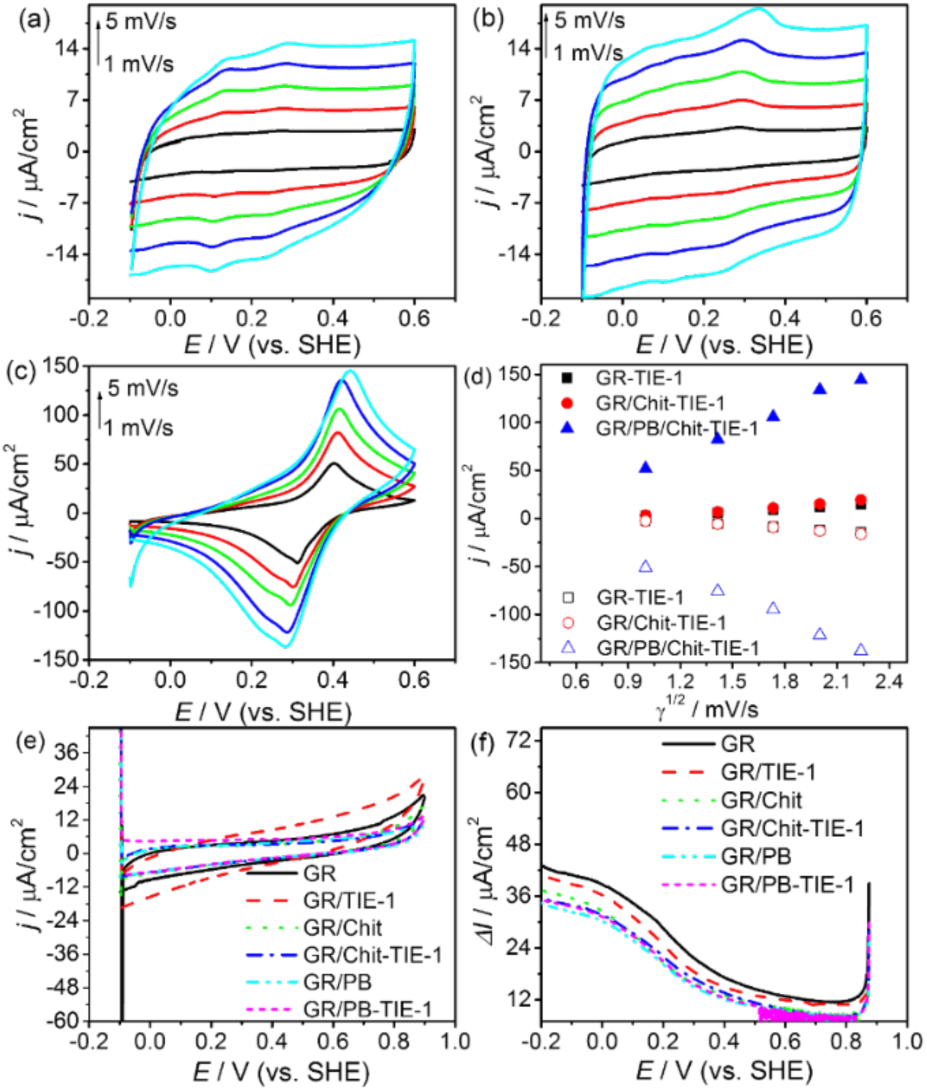
Scan rate dependence cyclic voltammetry of TIE-1 on graphite electrodes in 50 mM PBS (pH7); unmodified biocathode (a), biocathode modified with chitosan (b), biocathode modified with chitosan - Prussian blue (c); Linear relationship of anodic (solid symbols) and cathodic (open symbols) peak current with square root of scan rate, γ^1/2^ (d). Cyclic Voltammetry (e) at a scan rate of 5 mV/s and Differential Pulse Voltammetry (f) of cell-free spent medium (supernatant) at the end of EU experiment using a glassy carbon electrode.

### 3.6. Electrochemical Impedance spectroscopy (EIS) and potentiodynamic polarization of the biocathodes

EIS characterization of biocathodes was performed at the end of the EU experiment as shown in Fig. S3a. The EIS data were fitted into the equivalent circuit (EC) to derive the value of the circuit component [49,53]. The diameter of the semicircle corresponds to the charge transfer resistance (R_ct_) across the cathode/electrolyte interface. When the cathode interacts with microbes (biofilm), the R_ct_ value decreases gradually for all biocathodes compared to the corresponding negative control. The lower value of R_ct_ implies a faster bioelectrochemical reaction. Based on the simulated equivalent circuit, the R_ct_ value of the biocathodes was found to be 5145.2 ± 9.2 Ω (GR-TIE-1) >143.3 ± 2.2 Ω (GR/Chit-TIE-1) > 20.6 ± 2.8 Ω (GR/PB/Chit-TIE-1). The lower R_ct_ value might be due to the GR/PB/Chit-TIE-1 biocathode having an accelerated electrode reaction rate and higher current uptake as observed by CA and CV studies. Further, the lower R_ct_ of the modified biocathodes (e.g., Chitosan or PB complex cathode) can be explained by the nature of the ionically conductive biopolymer chitosan, which will help the bacterial cells make electrochemical contact with the electrode. The cathodes modified with an electron transfer mediator (PB complex) will enhance electron donation to bacteria, further lowering the R_ct_.

Potentiodynamic polarization (Tafel plots) of biocathodes was performed to evaluate the bioelectrochemical kinetics of surface bound redox probe, or PB modified cathodes with TIE-1 (Fig. S3b-c and Table S2). It indicates that the exchange current of GR/PB/Chit-TIE-1 was 10.3 ± 0.07 µA, which is about ten times higher than the unmodified biocathode (1.05 ± 0.02 µA, GR/TIE-1), and five times higher than the chitosan-based biocathode (1.9 ± 0.02 µA). The value of exchange current (*I*_*0*_) supports the current uptake trends observed in the EU and CV experiments. The biocathode potential at the intersection of the anodic and cathodic region for GR/PB/Chit-TIE-1 has a higher cathodic value (+45 mV) compared with the other biocathodes. The lower value of the anodic (β_*a*_ = 62.3 mV/dec) and cathodic (β_*c*_ = 197.2 mV/dec) slope can be attributed to the enhanced reaction rate of extracellular electron transfer at the biointerface of the GR/PB/Chit-TIE-1 biocathode.

Based on electrochemical analysis, the enhanced performance of PB based biocathodes is due to the reversible redox reaction between PB (Fe^3+^ site) and PW (Prussian White, Fe^2+^ site). At the cathodic reduction potential of +100 mV, the electrode surface bound with the Fe^2+^ complex (PW) is able to donate electrons continuously to TIE-1. Further, the microbially oxidized Fe^3+^ complex (PB) is cyclically reduced to Fe^2+^ complex (PW) by the poised potential enhancing extracellular electron transfer to TIE-1, and microbial electrosynthesis from CO_2_ (Fig. S1). From SEM images, it is evident that the attachment ability of TIE-1 clearly improved on the modified graphite cathode compared to the unmodified electrode (Fig. 9). Further, both modified cathodes consist of a network of chitosan, which appears to aid microbial attachment as supported by the higher current density utilized by TIE-1. The chitosan (biopolymer) is used to enhance the microbial attachment that we clearly see from SEM images of GR/Chit/TIE-1 (Fig. 9c). However, the electron uptake with and without chitosan shows similar values, which is likely due to the lack of redox active molecules in the chitosan. GR/PB/TIE-1 (no chitosan) was avoided due to potential issues of detachment/dissolution of PB without chitosan. The chitosan network holds the PB layer and provides an immobilized surface for microbial attachment (Fig. 9d). The “with and without chitosan” controls clearly show that microbial uptake is unaffected by the presence or absence of chitosan, further supporting the fact that chitosan does not affect microbial electron uptake significantly.

**Fig. 9.**
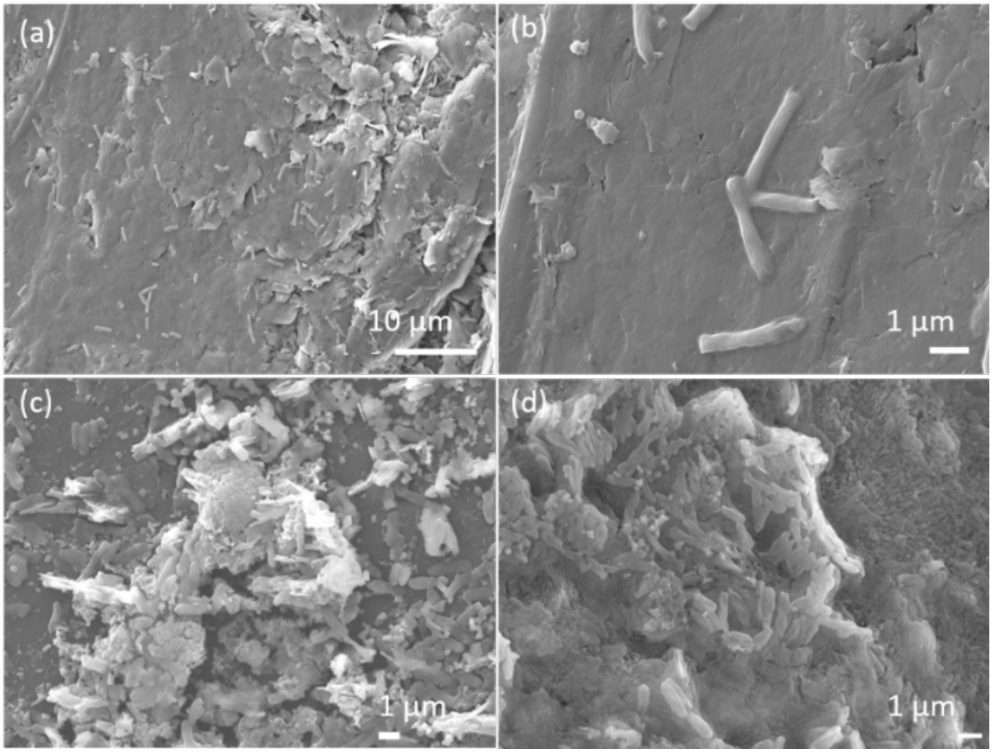
SEM images for the attachment of TIE-1 on different graphite electrodes, graphite (a, b), biocathodes modified with chitosan (c) and biocathodes modified with PB-Chitosan (d).

### 3.7. Implications on future MES studies

This work emphasizes that the PB modified graphite electrodes enhance direct electron transfer by *3.8 – fold* with respect to current density (0.0568 ± 0.09 A/m^2^) when compared to unmodified graphite. However, this effect is lost when we add Fe(II) to the system (Table 1). Further, the dissolved Fe(II) added to the medium is electrochemically and/or biologically oxidized to Ferrihydrite (oxides of Fe(III)) at the surface of the electrode as well as on the TIE-1 cell surface (Fig. 2, Fig. 3, Fig. 4 & Fig. S2). This oxidation was supported by electrochemical data (Table 1). Biotic reactors with Fe(II) showed anodic current in contrast to those coated with PB-Chitosan (GR/PB/Chit-TIE-1).

**Table 1.**
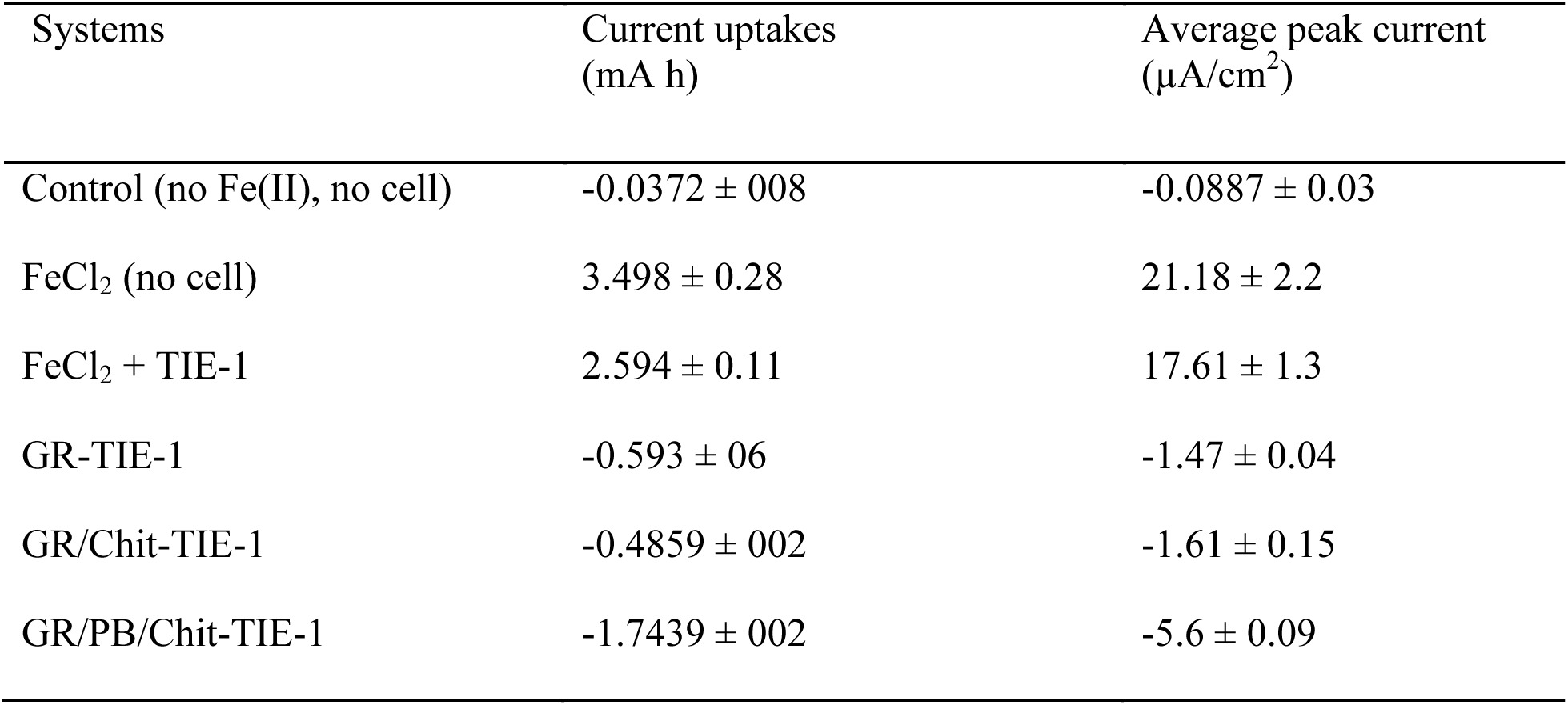
Summary of current consumption (n=3) with different systems at poised potential of +100 mV vs. SHE

Recently many researchers have explored the importance of direct electron uptake and utilization of electrons from various biocathodes in MESs for biofuel production [3,54,55]. MESs mimic the process of natural autotrophy by using carbon dioxide as a carbon source for biosynthesis [55]. In MES applications, a surplus amount of electron uptake is required to reduce carbon dioxide to biofuels/biochemical in contrast to the utilization of already reduced carbon sources (eg., sugars, glycerol) [55]. Hence, the high demand of electrons in MES can limit the production of biochemicals. Our modified biocathode with TIE-1 (GR/PB/Chit-TIE-1) showed a reproducible increase in electron uptake (*3.2-fold* higher for current consumption, −0.593 ± 06 mA h to −1.74 ± 0.03 mA h, and *3.8-fold* higher current density 1.47 ± 0.04 to 5.6 ± 0.09 µA/cm^2^). This effect can play a significant role in direct electron transfer strategies (biocathode poised at which no H_2_ production) in the field of MES [56]. For context, in a recent study authors showed that changing the electrode material to graphite felt and increasing the time of operation of a BEC with *Clostridium pasteurianum* increased both current density (−1.5 ~ −5 mA or −14 µA/cm^2^ ~ −46 µA/cm^2^, *3-fold* increase) and biobutanol production from glucose +45 mV vs. SHE (*6-fold* increase) [55]. This improvement in current density is in the range of what we report here for direct electron uptake by TIE-1 using a PB modified electrode. The increase in current density also led to increased biomass (*2-fold* higher) and PHB (*1.4-fold* higher), which is the first step toward improving bioproduction using TIE-1. Future work will explore the use of natural iron oxides coated electrodes as potential redox mediators for TIE-1.

## 4. Conclusions

In summary, electrodes modified with the redox complex Prussian blue (PB) improved electron transfer to the photoelectroautotroph, *Rhodopseudomonas palustris* TIE-1. The PB complex based biocathode showed increased cathodic current density (5.6 ± 0.09 µA/cm^2^), which is 3.8 times higher than the unmodified biocathode. A higher current uptake capacity (-1.744 ± 0.03 mA h for 130 h), enhanced production of the bioplastic, polyhydroxybutyrate (18.8 ±0.3 g/L, PHB), and lower charge transfer resistance (R_ct_, 20.6 ± 2.8 Ω) of the PB based biocathode suggests that the reversible redox nature of the PB complex acts as an electron transfer (ET) agent. Our results indicate that the modified biocathode offers an advantage to TIE-1 grown under photoelectroautotrophic conditions by increasing electron transfer rates and current density. TIE-1 is a prime candidate for microbial electrosynthesis, and these modified electrodes will aid higher bio-production of value-added biochemicals.

## Acknowledgements

The authors would like to acknowledge financial support from US Department of Energy (grant number DESC0014613) and the David and Lucile Packard Foundation to carry out this research. We also thank Mr. Michael Guzman, Washington University, USA for valuable comments.

---

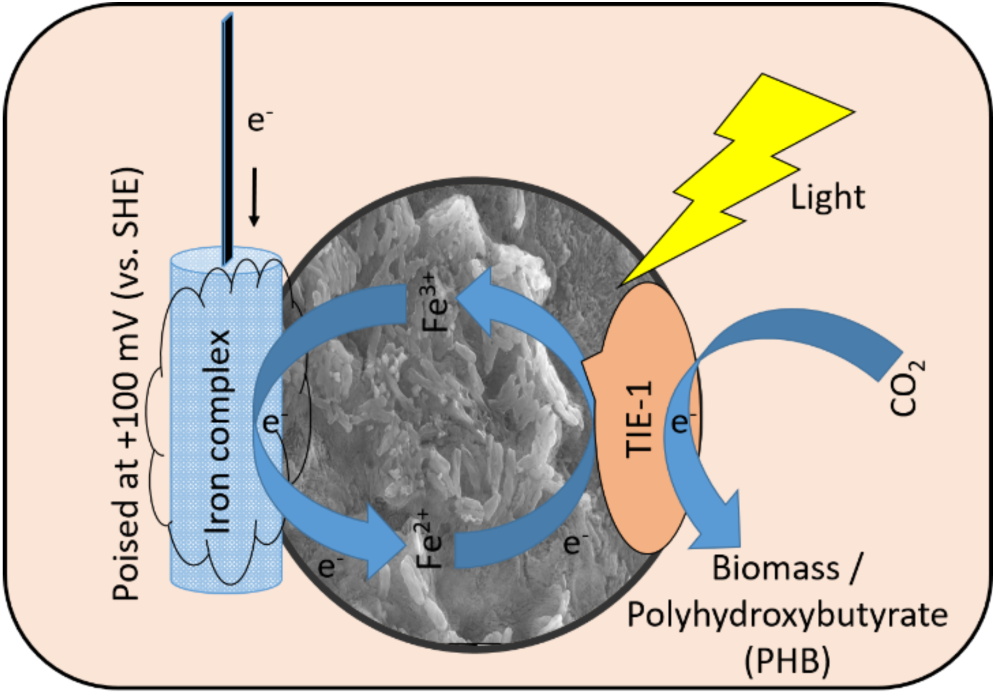
Graphical abstract / TOC:

